# IMPAIRED BRIDGING OF TEMPORAL DISCONTINUITIES IN OLDER ADULT HIV-1 TG RATS

**DOI:** 10.64898/2026.04.06.716768

**Authors:** Kristen A. McLaurin, Hailong Li, Allison Ritchie, Rosemarie M. Booze, Charles F. Mactutus

**Affiliations:** Program in Behavioral Neuroscience, Department of Psychology, Barnwell College, University of South Carolina, 1512 Pendleton Street, Columbia, SC 29208; Department of Pharmaceutical Sciences, College of Pharmacy, University of Kentucky, 789 South Limestone Street, Lexington, KY 40508

**Author notes:** **Address proofs and correspondence to:** Charles F. Mactutus, Ph.D., Department of Psychology, 1512 Pendleton Street, University of South Carolina, Columbia, SC 29208.

**Keywords:** Neurocognition, Attention, Dendritic Spines, Amyloid Beta

## Abstract

The advent and widespread uptake of combination antiretroviral therapy dramatically changed the epidemiological features of human immunodeficiency virus type 1 (HIV-1), whereby older individuals (>50 years of age) account for approximately 50% of HIV-1 seropositive individuals in the United States. Nevertheless, to date, there is no extant *in vivo* biological system to model the unique age-related neurocognitive impairments observed in HIV-1 seropositive individuals. Herein, the utility of the HIV-1 transgenic (Tg) rat as a biological system to model age-related neurocognitive impairments and neuroanatomical alterations was evaluated. Older adult HIV-1 Tg rodents (i.e., >12 months of age upon testing initiation), relative to their control counterparts, exhibited profound neurocognitive alterations characterized by impairments in stimulus-reinforcement learning, sustained attention, and selective attention; neurocognitive deficits which support a fundamental distortion of temporal processing. Neuronal dysfunction in older adult HIV-1 Tg animals was characterized by structural alterations in pyramidal neurons, and their associated dendritic spines, in the medial prefrontal cortex and abnormal accumulation of amyloid beta (Aβ). Interestingly, the abnormal accumulation of Aβ mechanistically underlies, at least in part, the profound dendritic spine dysmorphology in male, but not female, HIV-1 Tg rats. More critically, however, neuronal dysfunction mechanistically underlies neurocognitive impairments in both male and female HIV-1 Tg rodents, whereby neuronal dysfunction accounts for 65.4% and 60.8% of the variance in neurocognitive function, respectively. Establishing the utility of the HIV-1 Tg rat for age-related neurocognitive impairments is fundamental to disentangling the role of HIV-1 viral proteins and comorbidities in neurocognitive function.

## INTRODUCTION

The widespread uptake of combination antiretroviral therapy (cART; i.e., approximately 77% of individuals living with human immunodeficiency virus type 1 (HIV-1) were accessing cART at the end of 2024; (1)) dramatically shifted the demographic features of the disease. Indeed, prior to the advent of cART, a HIV-1 diagnosis was terminal; current life-expectancy estimates for HIV-1 seropositive individuals, in sharp contrast, resemble those for seronegative persons (2). As a result, older individuals (>50 years of age) account for approximately 25% of HIV-1 seropositive individuals globally (3) and over 50% of individuals with HIV-1 in the United States (4). Therefore, understanding the unique age-related health challenges, including an increased risk for neurocognitive impairments, faced by older HIV-1 seropositive individuals, is of utmost concern.

In 2007, the research nosology guiding the diagnosis of neurologic manifestations of HIV-1 was updated to a primarily neurocognitive disorder collectively termed HIV-1-associated neurocognitive disorders (HAND; (5)). Among HIV-1 seropositive individuals, the overall prevalence of HAND ranges from 24% to 50% (6–7); older age, in particular, is considered a risk factor for HAND (6–7). Indeed, in cross-sectional studies, older HIV-1 seropositive individuals exhibit a higher frequency of neurocognitive impairments relative to their younger counterparts (8–11) and non-infected older adults (8). More specifically, older HIV-1-infected individuals, relative to seronegative adults, display poorer performance in the neurocognitive domains of processing speed, verbal, recall, motor/psychomotor, and executive function (10). Nevertheless, despite strong evidence for the fundamental role of age in the prevalence of HAND, methodological limitations (e.g., failure to include non-infected controls and/or control for comorbidities; (12)) limit our understanding of how aging impacts neurocognitive function in HIV-1 seropositive individuals.

An extant *in vivo* biological system to model the unique age-related neurocognitive impairments observed in HIV-1 seropositive individuals affords a fundamental opportunity to address this limitation. In 2001, Reid *et al.* (13) developed a non-infectious HIV-1 transgenic (Tg) rat consisting of a *gag-pol* deleted HIV-1 provirus regulated by the human long terminal repeat. Expression of HIV-1 viral proteins constitutively throughout development in the HIV-1 Tg rat (14–15) resembles HIV-1 seropositive individuals on cART and heralds an opportunity to evaluate the consequences of long-term HIV-1 viral protein exposure. The current HIV-1 Tg rat derivation, with the transgene now limited to chromosome 9 and in the F344/N background strain, is healthier (i.e., Similar Growth Rates (14); Longer Lifespan (14); Intact Sensory and Gross Motoric System Function Through Advancing Age (16)) than those originally described. Indeed, longitudinal studies in the HIV-1 Tg rat have been instrumental in defining HAND as a neurodegenerative disease characterized by age-related disease progression (16–17); findings which are consistent with time-limited studies in clinical populations, whereby a significant proportion (i.e., 13-30%) of HIV-1 seropositive individuals exhibit overall neurocognitive decline across time (e.g., 18-19). Despite the undeniable strength of the prior longitudinal studies in the HIV-1 Tg rat, the true impact of long-term viral protein exposure on neurocognitive decline was potentially minimized by the process of repeated testing; a process that affords an opportunity to maintain a cognitive reserve.

To address this knowledge gap, a longitudinal experimental design without the confound a repeated testing was implemented to evaluate the true magnitude of neurocognitive and neuroanatomical decline in the HIV-1 Tg rat. Male and female older adult (>12 months of age) F344/N HIV-1 Tg and control rodents were evaluated in a series of operant tasks tapping higher-order cognitive processes, including stimulus-reinforcement learning, sustained attention, selective attention, discrimination, and extradimensional set-shifting. Structural alterations to pyramidal neurons in layers II-III of the medial prefrontal cortex (mPFC) and amyloid-β (Aβ) protein levels were examined after the completion of neurocognitive testing to evaluate neuronal dysfunction. Exploratory multiple regression analyses were conducted to establish neuronal dysfunction as a key neural mechanism underlying neurocognitive impairments in older adult rodents. Establishing the utility of the HIV-1 Tg rat for age-related neurocognitive impairments is fundamental to disentangling the role of HIV-1 viral proteins and comorbidities in neurocognitive function.

## MATERIALS AND METHODS

### Animals

Fischer F344/N control rats (Male: *n*=14; Female: *n*=23) were procured from Inotiv Laboratories Inc. (Indianapolis, IN), whereas Fischer F344/N HIV-1 Tg rodents were bred (Female Fischer F344/N Control x Male Fischer F344/N HIV-1 Tg; Male: *n*=42; Female: *n*=24) at the University of South Carolina (USC). Rodents were at least 12 months of age when neurocognitive testing was initiated. Due to health issues (e.g., significant weight loss, tumors), some animals were euthanized prior to the completion of at least a subset of assessments (i.e., Visual Prepulse Inhibition, Locomotor Activity, Completion of the Second Vigilance Program), yielding a subset of animals for analysis; sample sizes are described for each neurocognitive domain below or in Supplementary Information.

Animals were pair- or group-housed with animals of the same sex for the duration of the experiment. Rodents had *ad libitum* access to chow and water and were maintained in AAALAC-accredited facilities at the USC using guidelines established by the National Institute of Health. Environmental conditions for the animal colony were targeted at: 21°± 2°C, 50% ± 10% relative humidity and a 12-h light:12-h dark cycle with lights on at 0700 h (EST). The USC Institutional Animal Care and Use Committee approved the project protocol under federal assurance (#D16-00028).

### Integrity of Visual System Function: Visual Prepulse Inhibition

#### Apparatus

The startle platform (San Diego Instruments, Inc., San Diego, CA), a high-frequency loudspeaker (RadioShack Model #40–1278B), and a white LED light (22 lux) were housed within an 81 x 81 x 116 cm (External Dimensions) double-walled isolation cabinet (Thickness: 10 cm; Industrial Acoustic Company, Inc., Bronx, NY); 30 dB(A) of sound attenuation was provided within the chamber relative to the external environment. A high-frequency loudspeaker, which was mounted 30 cm above the Plexiglas animal enclosure, delivered the auditory prepulse (85 db(A), 20 msec duration) and startle stimuli (100 db(A), 20 msec duration). A white LED light (22 lux) was hung on the wall in front of the test cylinder for the presentation of a 20 msec visual prepulse stimuli. The animal’s ballistic response to the startle stimuli induced a deflection of the Plexiglas animal enclosure; the deflection was converted into analog signals by the piezoelectric accelerometer attached to the bottom of the test cylinder. Analog signals were recorded (2000 samples per sec), digitized (12-bit A to D) and saved to a hard disk.

#### Procedure

Older adult rodents were habituated and assessed in cross-modal prepulse inhibition (PPI) to evaluate the integrity of visual system function. Additional methodological details are presented in the Supplementary Information Text available online.

### Integrity of Gross Motoric System Function: Locomotor Activity

#### Apparatus

Perspex inserts were utilized to convert square (40 x 40 cm) activity monitors (Hamilton Kinder, San Diego Instruments, San Diego, CA) into round compartments (∼40 cm diameter). Gross movement was detected using infrared photocell interruptions; the activity monitors contained 32 emitter/detector pairs. The manufacturer tuned the photocells to maintain their sensitivity with the additional layer of perspex.

#### Procedure

Three 60-min locomotor activity assessments were conducted on consecutive days to evaluate gross motoric system function in older adult HIV-1 Tg (Male: *n*=10; Female: *n*=17) and control animals (Male: *n*=12; Female: *n*=16). Assessments were conducted in an isolated room between 700 and 1200h (EST) under dim light conditions. The number of photocell interruptions within the test session were collected for analysis.

### Components of Executive Function

#### Apparatus

Rodents were evaluated in a series of neurocognitive tasks using operant boxes housed inside sound-attenuating chambers (Med Associates, Inc., Fairfax, VT). The front wall of the operant box contained a 45 mg pellet dispenser stationed between two retractable levers and three panel lights; one panel light was positioned above each lever (Incandescent, 22 lux), and one panel light was stationed above the pellet dispenser (LED, 11 lux). An incandescent house light was located at the top of the rear wall of the operant chamber. Signal presentation, lever operation, reinforcement delivery, and data collection were controlled by a personal computer (PC) and Med-PC for Windows software (V 4.1.3; Med Associates).

#### Sustained Attention: Signal Detection Operant Task

Older adult rodents were initially trained to lever-press for sucrose pellet rewards using a standard shaping response protocol across approximately 35 sessions. Rodents were subsequently trained in a signal detection operant task originally described by McGaughy and Sarter (20). The signal detection operant task utilizes a series of three vigilance programs to train animals to discriminate between the presence and absence of a visual stimulus (i.e., central panel light illumination and no illumination, respectively) across 160-162 testing trials. The third vigilance program, during which the length of the visual stimulus was systematically manipulated (1000, 500, 100 msec), is of particular import to the present study. Additional methodological details are available in the Supplementary Information Text available online and described in detail by McLaurin et al. (21).

#### Selective Attention: Visual Distractor Task

After successfully acquiring the signal detection operant task, selective attention was evaluated in HIV-1 Tg (Male: *n*=6; Female: *n*=17) and control (Male: *n*=12; Female: *n*=12) animals for five test sessions using a visual distractor task. The third vigilance program of the signal detection operant task was divided into three trial blocks, whereby each trial block contained 54 trials. During the second trial block, a 1.5 sec visual distractor stimulus (i.e., the house light) was presented 1 sec prior to and 0.5 sec after the stimulus (i.e., central panel light illumination) onset. Statistical analyses and figures represent all five days in the task.

#### Stimulus Discrimination

Subsequently, the cognitive domain of stimulus discrimination was evaluated in older adult rodents using a discrimination operant task (22). In brief, 1 sec left or right panel visual stimulus trials were presented randomly across a 160-trial session, whereby animals were rewarded with a sucrose pellet for responding on the lever underneath the light stimulus. For successful acquisition of the discrimination operant task, rodents were required to respond at least five times and achieve 65% accuracy, calculated as (((Number of Correct Responses)/(Total Number of Responses)) x 100) for three consecutive or five non-consecutive days. Statistical analyses and figures represent the days meeting criteria during the discrimination task, yielding samples sizes of: (Control Male: *n*=12; Control Female: *n*=11; HIV-1 Tg Male: *n*=1; HIV-1 Tg Female: *n*=17).

#### Extradimensional Set-Shifting

Lastly, older adult HIV-1 Tg (Male: *n*=1; Female: *n*=17) and control animals (Male: *n*=12; Female: *n*=11) were tested in an extradimensional set-shifting task (22). In brief, 1 sec left and right visual stimulus trials were randomly presented across a 160-trial session. Herein, the presence or absence of a visual stimulus indicated which lever to press, with half of the animals being rewarded for responding on the left lever and half of the rodents being rewarded for pressing the right lever. Rodents were required to meet the criterion defined for discrimination to successfully acquire the extradimensional set-shifting task. Statistical analyses and figures represent the days meeting criteria during the discrimination task.

### Post-mortem Analysis

#### Rodent Tissue Preparation

Rodents were anesthetized (5% Sevofluorane; Abbot Laboratories, North Chicago, IL) and transcardially perfused with 4% paraformaldehyde. After perfusion, the rodent brain was post-fixed in 4% paraformaldehyde for 10 minutes and sliced coronally using a rat brain matrix (500 μm; ASI Instruments, Warren, MI). Coronal brain slices were obtained from the mPFC (located approximately 3.7 mm to 2.2 mm anterior to Bregma; (23))

#### Synaptodendritic Dysfunction: Ballistic Labeling

Methodology for the ballistic labeling technique used to evaluate synaptodendritic alterations in pyramidal neurons in the mPFC has been previously reported in detail by Li et al. (24). In brief, ballistic cartridges were prepared and a Helios gene gun (Bio-Rad, Hercules, CA) was used to fire the tungsten bead/DiIC19(3) bead tubing onto the brain slices. Z-stack images (60x Oil Objective, *n.a.*=1.4, Z-Plane: 0.15 μm) of pyramidal neurons in the mPFC were acquired using a Nikon TE-2000E confocal microscopy system in combination with Nikon’s EZ-C1 software (Version 3.81b). Pyramidal neurons, and their associated dendritic spines, from the mPFC were analyzed using Neurolucida 360 (MicroBrightfield, Williston, VT).

#### Amyloid-β: Immunohistochemistry

Coronally sectioned rodent brain tissues (Control Male: *n*=12; Control Female: *n*=11; HIV-1 Tg Male: *n*=7; HIV-1 Tg Female: *n*=18) were incubated overnight at 4°C with a fluorescently labeled primary antibody for Aβ (i.e., Alexa Fluor® 488 Anti-β-Amyloid 1-42 Antibody, Catalogue Number ab224026, Abcam, Waltham, MA). Brain sections were mounted and fluorescent images were acquired using a Nikon TE-2000E confocal microscope system. ImageJ software was utilized to quantify the fluorescent signals.

### Statistical Analysis

Data were analyzed using analysis of variance (ANOVA) and regression techniques (SAS/STAT Software 9.4, SAS Institute, Inc., Cary, NC; SPSS Statistics 31, IBM Corp., Somer, NY; GraphPad Prism 10, GraphPad Software, Inc., La Jolla, CA). Figures were created using GraphPad Prism 10. The sphericity assumption in repeated-measures ANOVA analyses was accounted for via either the conservative Greenhouse-Geisser *df* correction factor (*p*_GG_) or orthogonal decomposition of the repeated-measures factors. Statistical significance was established at an alpha value of *p* ≤ 0.05. The Supplementary Information Text contains additional details on the statistical analysis.

## RESULTS

### Visual system function and gross motoric system function remain intact in older adult HIV-1 Tg and control rats

Visual PPI was conducted to establish the integrity of visual system function in older adult HIV-1 Tg and control animals. In PPI, the presentation of a punctate prestimulus 30-500 msec prior to the startling stimulus induces a dramatic decrease in auditory startle response (16). Indeed, independent of genotype, rodents exhibited robust inhibition to the visual prepulse at the 50 msec interstimulus interval (ISI; Supplementary Figure 1A; Main Effect: ISI, [*F*(5,255)=30.4, *p*_GG_≤0.001, η_p_ ^2^=0.374] with a prominent quadratic component [*F*(1,51)=45.2, *p*≤0.001, η_p_^2^=0.470]).

The integrity of gross motoric system function was confirmed via the mean number of total photocell interruptions across three 60-min locomotor activity test sessions. Independent of biological sex, HIV-1 Tg animals exhibited a significantly fewer mean number of photocell interruptions relative to control animals (Supplementary Figure 1B; Main Effect: Genotype [*F*(1,51)=26.3, *p*≤0.001, η_p_^2^=0.340]). Complementary one-sample *t*-tests were conducted independently for each genotype to statistically test whether the average number of photocell interruptions was significantly greater than 0 (HIV-1 Tg: [*t*(26)=26.2, *p*≤0.001]; Control [*t*(27)=33.4, *p*≤0.001]). Therefore, consistent with previous studies through advanced age (16), there is strong evidence for the integrity of visual and gross-motoric system function in older adult HIV-1 Tg and control animals; the HIV-1 Tg rat, therefore, is a valid biological system to model neurocognitive impairments in older HIV-1 seropositive individuals on cART.

### Older adult HIV-1 Tg rodents, independent of sex, exhibited selective neurocognitive impairments relative to control animals

#### Sustained Attention: Signal Detection Operant Task

Rodents successfully acquired the signal detection operant task, comprised of three vigilance programs, upon meeting the criterion of 65% accuracy for three consecutive or five non-consecutive test sessions. The third vigilance program systematically manipulated the signal duration to critically evaluate the temporal component of sustained attention. Indeed, prominent impairments in stimulus-reinforcement learning (Figure 1A), sustained attention (Figure 1B), and the temporal component of sustained attention (Figure 1C) were observed in HIV-1 Tg, relative to control, rats in the third vigilance program of the signal detection operant task.

**Figure 1.**
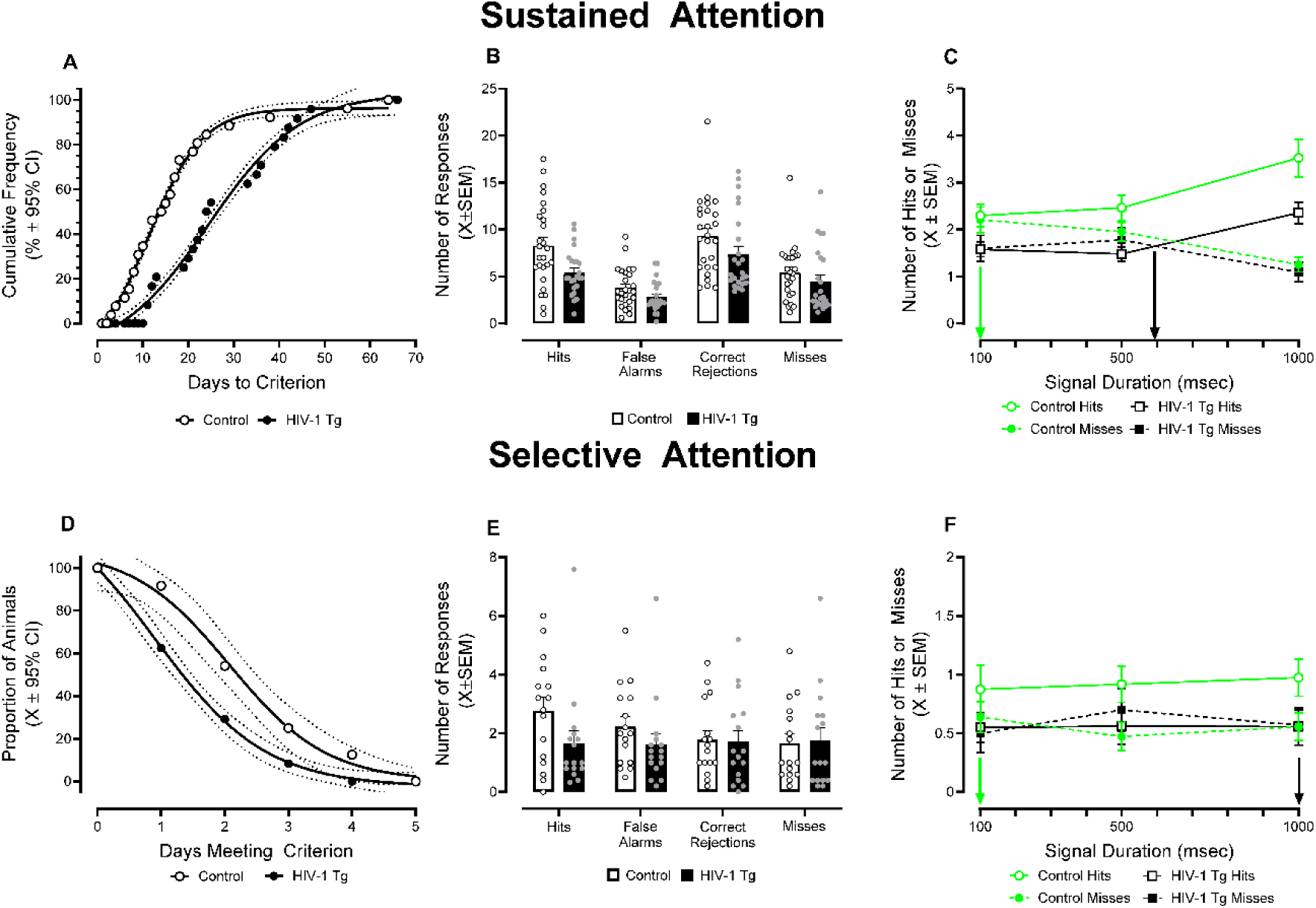
Constitutive Expression of HIV-1 Viral Proteins Impairs Sustained and Selective Attention. A signal detection operant task was conducted to evaluate key aspects of sustained attention (A-C) and selective attention (D-F). **(A)** Rodents successfully acquired the third vigilance program of the signal detection operant task by meeting the criterion of 65% accuracy for three consecutive or five non-consecutive test sessions. HIV-1 transgenic (Tg) rats, independent of biological sex, exhibited a profound impairment in stimulus-reinforcement learning, evidenced by a slower rate of task acquisition relative to their control counterparts. **(B)** During the third vigilance program of the signal detection operant task, a rodent may emit one of four responses, including hits, false alarms, correct rejections, and misses. HIV-1 Tg rodents exhibited significantly fewer hits relative to their control counterparts. **(C)** The length of the stimulus signal was systematically manipulated to evaluate the temporal components of sustained attention. HIV-1 Tg animals, relative to controls, exhibited a profound rightward shift in the loss of signal detection (i.e., the time at which the number of hits and misses intersect indicated by downward arrows pointing at the x-axis). **(D)** Selective attention was evaluated for five test sessions by presenting a 1.5 sec visual distractor at the beginning of Trials 54 to 108 in the third vigilance program of the signal detection operant task. HIV-1 Tg rodents met criteria (i.e., 65% accuracy) on significantly fewer days than their control counterparts. **(E)** The number of hits, false alarms, correct rejections, and misses was evaluated during Trials 54 to 108. In the presence of a visual distractor, HIV-1 Tg rats failed to successfully acquire the task contingencies. **(F)** When the length of the stimulus signal was systematically manipulated during Trials 54 to 108, HIV-1 Tg rodents failed to accurately detect the signal, evidenced by a loss of signal detection at 1000 msec.

Examination of the number of days to meet criteria in the third vigilance program revealed a profound HIV-1-transgene-induced deficit in stimulus-reinforcement learning. A sigmoidal dose-response curve with a variable slope afforded the best-fit function for task acquisition in both control (*R*^2^=0.99) and HIV-1 Tg (*R*^2^=0.98) rats, albeit significant differences in the fit of the function were observed (Figure 1A; [*F*(4,42)=154.3, *p*≤0.001]). HIV-1 Tg rodents exhibited an initial eight-day lag before any of the HIV-1 Tg animals met the criterion, as well as a slower rate of task acquisition; the significantly slower rate of task acquisition is confirmed via an evaluation of differences in the Hill Slope parameter of the sigmoidal dose-response curve with a variable slope [*F*(1,42)=9.8, *p*≤0.001].

Presence of the HIV-1 transgene also significantly influenced the rodent’s response profile (i.e., Number of Hits, False Alarms, Correct Rejections, Misses) during the third vigilance program of the signal detection operant task (Figure 1B; Genotype x Response Type Interaction, [*F*(3,138)=28.6, *p*≤0.001]). Multiple comparisons were conducted on the interaction revealing fewer hits in HIV-1 Tg rodents relative to controls [*t*(138)=3.22, *p*≤0.033). The factor of biological sex influenced the magnitude, but not direction, of the interaction (Genotype x Sex x Response Type Interaction, [*F*(6,138)=23.5, *p*≤0.001]).

Further, independent of biological sex, HIV-1 Tg rodents exhibited a profound impairment in the temporal components of sustained attention across varying signal durations (Figure 1C; Genotype x Response Type x Duration Interaction, [*F*(2,92)=4.3, *p*≤0.017]). Specifically, HIV-1 Tg animals exhibited a prominent rightward shift in the loss of signal detection (i.e., the time at which the number of hits and misses intersect) relative to control rodents (approximately 595 msec vs 100 msec, respectively).

#### Selective Attention: Visual Distractor Task

Subsequently, the neurocognitive domain of selective attention was evaluated for five test sessions, whereby a 1.5 sec visual distractor was presented at the beginning of each trial in the second trial block (i.e., Trials 54-108). A visual distractor task revealed profound deficits in HIV-1 Tg rodents, relative to control animals, in the neurocognitive domains of stimulus-reinforcement learning (Figure 1D), selective attention (Figure 1E), and the temporal components of selective attention (Figure 1F).

Examination of the number of days rodents met criteria in the visual distractor task revealed a profound impairment in stimulus-reinforcement learning in HIV-1 Tg animals. The number of days HIV-1 Tg and control rats met criteria in the visual distractor task was well-described by a sigmoidal dose-response curve with a variable slope (Control: *R*^2^=0.99; HIV-1 Tg: *R*^2^=0.99). Nevertheless, genotype significantly influenced the fit of the function [*F*(4,4)=15.2, *p*≤0.011], whereby a prominent leftward shift (i.e., Statistically Significant Difference in the LogEC_50_ [*F*(1,4)=10.1, *p*≤0.034]) in the fit of the function was observed in HIV-1 Tg animals relative to control rodents. Indeed, the vast majority (i.e., 70.8%) of HIV-1 Tg animals either failed to meet criteria during any test session or met criteria during only one test session.

Presence of the HIV-1 transgene also significantly influenced the rodent’s response profile during the second trial block of the visual distractor task (Figure 1E; Genotype x Response Type Interaction, [*F*(3,90)=8.7, *p*≤0.001]). Complementary analyses were conducted by genotype to elucidate the locus of the interaction, whereby control (Main Effect: Response Type, [*F*(3,45)=19.5, *p*≤0.001]), but not HIV-1 Tg (Main Effect: Response Type, *p*>0.05) , successfully acquired the task contingencies during the second trial block.

Examination of responses during signal trials (i.e., Hits, Misses) across varying signal durations confirmed these findings (Figure 1F; Genotype x Response Type Interaction, [*F*(1,30)=8.6, *p*≤0.001]). Multiple comparisons, which were conducted on the interaction, revealed that control [*t*(30)=3.8, *p*≤0.004], but not HIV-1 Tg (*p*>0.05), rats differentiated between hits and misses. Indeed, HIV-1 Tg animals exhibited a prominent rightward shift in the loss of signal detection relative to control rodents (approximately 1000 msec vs 100 msec, respectively).

#### Stimulus Discrimination

Older adult HIV-1 Tg and control animals were next evaluated in a visual discrimination task, which taps the rodent’s ability to distinguish between stimuli. Selective impairments in stimulus discrimination were observed in HIV-1 Tg relative to control animals (Stimulus-Reinforcement Learning: Figure 2A; Visual Discrimination: Figure 2B).

**Figure 2.**
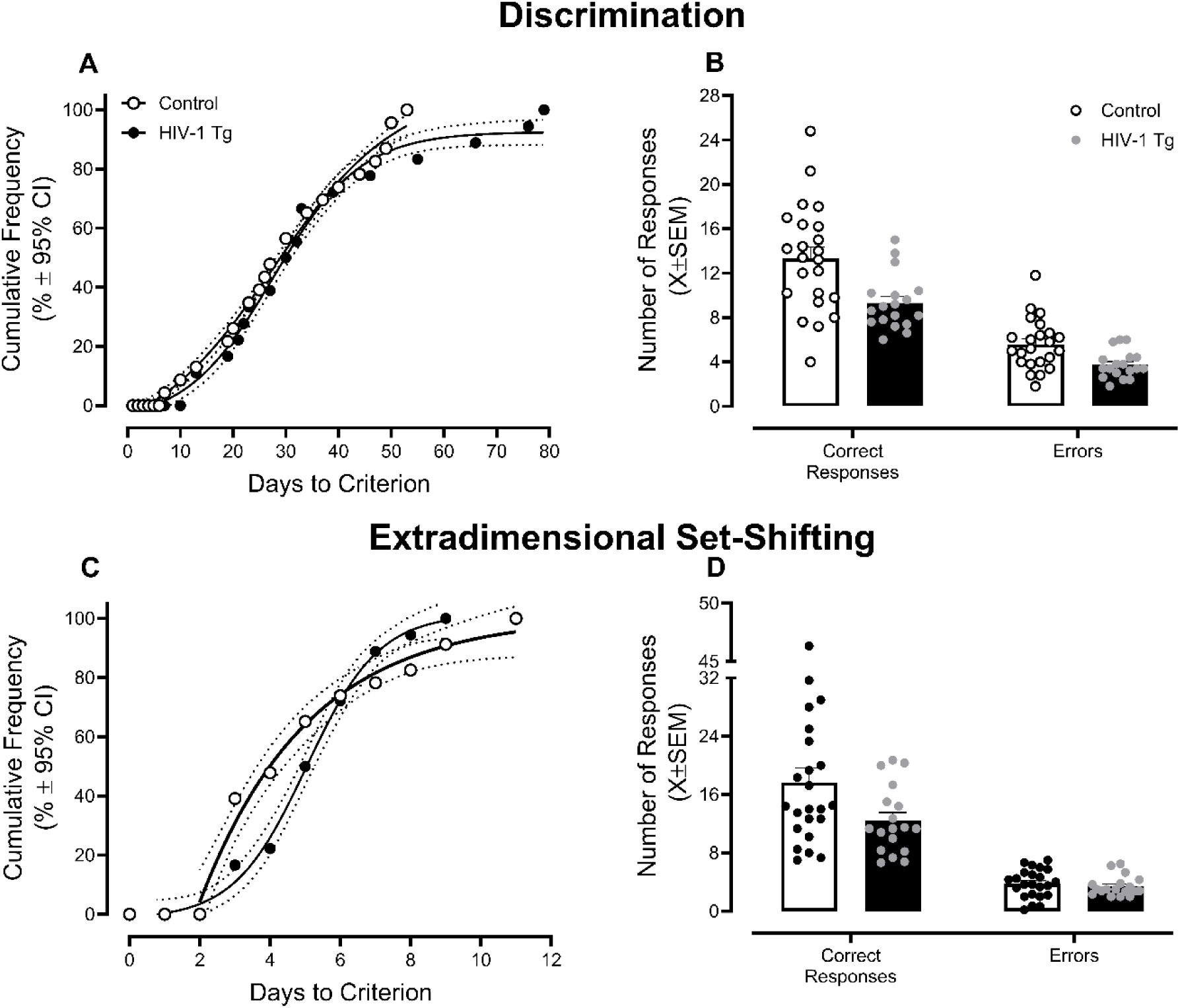
HIV-1 Transgenic (Tg) Rats Exhibited Selective Alterations in Discrimination and Extradimensional Set-Shifting. An operant task was utilized to evaluate the rodent’s ability to distinguish between stimuli (i.e., Discrimination) or shift their attention from one dimension to another (i.e., Extradimensional Set-Shifting). **(A)** Rodents successfully acquired the discrimination task by meeting the criterion of 65% accuracy for three consecutive or five non-consecutive sessions. HIV-1 Tg rats exhibited a profound impairment in stimulus-reinforcement learning, evidenced by the approximately 20% of animals that required over 53 days to meet criterion. **(B)** During the discrimination task, a rodent may emit one of two responses, including a correct response or an error. Although HIV-1 Tg animals exhibited significantly fewer overall responses, there was no compelling evidence for impairments in stimulus discrimination. **(C)** In the cognitive domain of extradimensional set-shifting, older adult HIV-1 Tg rodents exhibited a significant impairment in stimulus-reinforcement learning, evidenced by requiring a greater number of days to meet criterion than their control counterparts. **(D)** Older adult HIV-1 Tg rats displayed no impairments in the extradimensional set-shifting task, as there were no statistically significant genotype differences in the number of correct responses or errors.

Presence of the HIV-1 transgene induced a statistically significant deficit in stimulus-reinforcement learning (Figure 2A). The number of days required for HIV-1 Tg and control rats to meet criteria in the visual discrimination task was well-described by a sigmoidal dose-response curve with a variable slope (Control: *R*^2^=0.99; HIV-1 Tg: *R*^2^=0.99), albeit statistically significantly differences in the fit of the function were observed [*F*(4,39)=3.6, *p*≤0.01]. Indeed, genotypic differences in the “top” parameter of the function [*F*(1,39)=5.9, *p*≤0.02] were observed reflecting the approximately 20% of HIV-1 Tg animals requiring over 53 days to meet criterion.

The number of correct and incorrect responses were subsequently examined for HIV-1 Tg and control animals. HIV-1 Tg, relative to control, rodents made significantly fewer overall responses (Main Effect: Genotype, [*F*(1,78)=17.0, *p*≤0.001]). The failure to observe a statistically significant interaction between genotype and response type (*p*>0.05), however, supports no compelling impairments in stimulus discrimination.

#### Extradimensional Set-Shifting

Lastly, rodents were evaluated in an extradimensional set-shifting task; an assessment that required animals to shift their attention from one dimension (i.e., location of the visual stimulus) to another (i.e., lever location). HIV-1 Tg animals, relative to their control counterparts, exhibited selective impairments in the extradimensional set-shifting task (Stimulus-Reinforcement Learning: Figure 2C; Extradimensional Set-Shifting: Figure 2D).

A prominent HIV-1-induced impairment in stimulus-reinforcement learning in the extradimensional set-shifting task was evidenced by an evaluation of the number of days to meet criterion (Figure 2C). Indeed, for control and HIV-1 Tg animals, the number of days to meet criterion in the extradimensional set-shifting task were well-described by a one-phase association (*R*^2^=0.98) and sigmoidal dose-response curve with a variable slope (*R*^2^=0.99), respectively. Differences in the best-fit function highlight the initial impairment exhibited by HIV-1 Tg rodents before any of the animals met criteria relative to the control rats.

Presence of the HIV-1 transgene, however, failed to alter the rodent’s response profile (i.e., correct and incorrect responses; Figure 2D; *p*>0.05).

### Morphological parameters of pyramidal neurons, and associated dendritic spines, in the medial prefrontal cortex were altered in a sex-dependent manner in older adult HIV-1 Tg rodents relative to their control counterparts

Post-mortem, neuronal dysfunction in the mPFC was evaluated using two complementary methods, including ballistic labeling of pyramidal neurons in layers II-III of the the mPFC and immunohistochemistry of Aβ.

#### Neuronal Morphology

The morphology of pyramidal neurons was evaluated using two complementary approaches, including the classic Sholl analysis (Figure 3A) and a centrifugal branch ordering scheme (Figure 3B-C), to examine neuronal arbor complexity and dendritic branching complexity, respectively.

**Figure 3.**
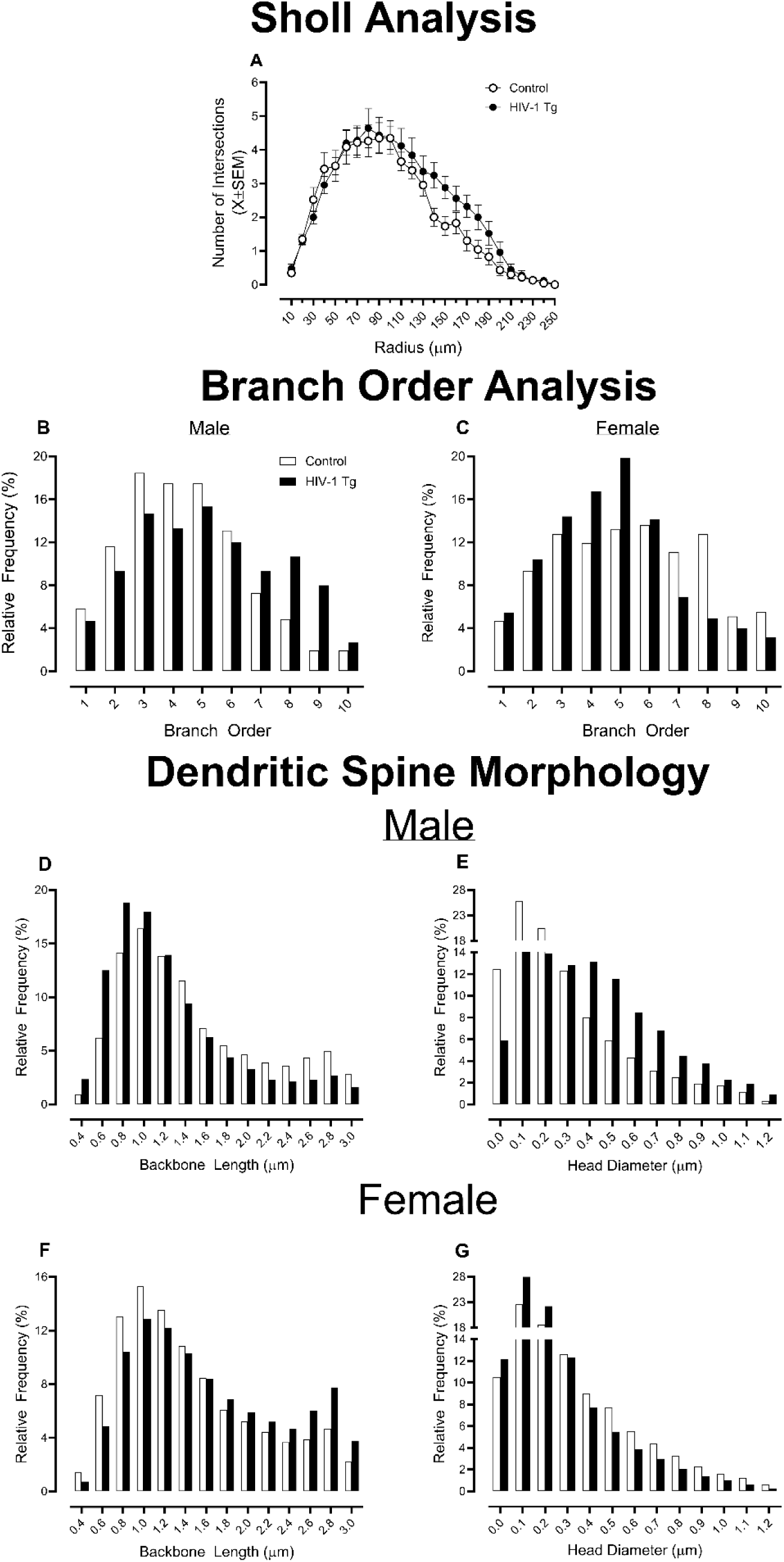
Presence of the HIV-1 Transgene Altered the Morphological Parameters of Pyramidal Neurons, and Their Associated Dendritic Spines, in the Medial Prefrontal Cortex. An innovative DiOlistic labeling technique was implemented to label pyramidal neurons in the medial prefrontal cortex of older adult HIV-1 transgenic (Tg) and control animals. **(A)** A classic Sholl analysis revealed that HIV-1 Tg rodents, independent of biological sex, had a greater number of dendritic intersections at higher radii relative to control animals; findings which support increased neuronal arbor complexity. **(B-C)** A complementary centrifugal branch ordering method was also implemented, whereby presence of the HIV-1 transgene significantly altered dendritic branching complexity in a sex-dependent manner. Indeed, HIV-1 Tg male rodents exhibited an increased relative frequency of dendritic branches at higher branch orders relative to their control counterparts. In sharp contrast, female HIV-1 Tg, relative to control, animals displayed an increased relative frequency of dendritic branches at lower branch orders. **(D-G)** Sophisticated neuronal reconstruction software was used to quantify two key morphological parameters of dendritic spines, including backbone length and head diameter; morphological parameters were altered in a genotype and sex dependent manner. HIV-1 Tg, relative to control, male rodents exhibited a population shift towards decreased dendritic spine backbone length and increased head diameter. In contrast, HIV-1 Tg female rats displayed a population shift towards increased dendritic spine backbone length and decreased head diameter relative to control animals.

In the Sholl analysis, the number of dendritic intersections occurring every 10 μm from the soma were quantified in older adult HIV-1 Tg and control animals. Independent of biological sex, HIV-1 Tg rodents exhibited increased neuronal arbor complexity relative to their control counterparts (Figure 3A; Genotype x Radius Interaction, [*F*(24,1056)=1.6, *p*≤0.03]). Specifically, when compared to control animals, HIV-1 Tg rats had a greater number of dendritic intersections at higher radii, supporting increased neuronal arbor complexity farther away from the soma.

A complementary centrifugal branch ordering method was also implemented to evaluate dendritic branching complexity, whereby each dendrite was assigned a branch order by counting the number of segments traversed. Presence of the HIV-1 transgene significantly altered dendritic branching complexity in a sex-dependent manner (Figure 3B-C; Genotype x Sex x Branch Order Interaction, [*F*(9,396)=31.8, *p*≤0.001]). Complementary analyses were conducted independently for male and female rodents to identify the locus of the interaction. Indeed, HIV-1 Tg male rodents exhibited an increased relative frequency of dendritic branches at higher branch orders relative to their control counterparts (Genotype x Branch Order Interaction, [*F*(9,153)=9.6, *p*≤0.001]). In sharp contrast, HIV-1 Tg, relative to control, female animals displayed an increased relative frequency of dendritic branches at lower branch orders (Genotype x Branch Order Interaction, [*F*(9,243)=31.0, *p*≤0.001]).

#### Dendritic Spine Morphology

Two fundamental morphological parameters of dendritic spines (i.e., Backbone Length (Figure 3D, 3F); Head Diameter (Figure 3E, 3G)), which serve as the primary postsynaptic compartment of excitatory synapses, were quantified using sophisticated neuronal reconstruction software. Both dendritic spine backbone length (Genotype x Sex x Bin Interaction, [*F*(18,792)=65.6, *p*≤0.001]) and head diameter (Genotype x Sex x Bin Interaction, [*F*(12,528)=118.6, *p*≤0.001]) were altered in a genotype- and sex-dependent manner.

To identify the locus of the interaction, complementary analyses were conducted independently by sex. Older adult male HIV-1 Tg rodents exhibited a population shift towards decreased dendritic spine backbone length (Genotype x Bin Interaction, [*F*(18,306)=37.8, *p*≤0.001]) and increased head diameter (Genotype x Bin Interaction, [*F*(12,204)=86.5, *p*≤0.001]) relative to control male rats. In sharp contrast, older adult HIV-1 Tg, relative to control, female animals displayed a population shift towards increased dendritic spine backbone length (Genotype x Bin Interaction, [*F*(18,486)=35.7, *p*≤0.001]) and decreased head diameter (Genotype x Bin Interaction, [*F*(12,324)=35.1, *p*≤0.001]). Relative to control animals, therefore, male and female HIV-1 Tg rodents exhibited a shift towards a stubby and thin dendritic spine phenotype.

#### Amyloid Beta

Aβ, an abnormal protein aggregate produced by the amyloidogenic processing of amyloid precursor protein, accumulates intraneuronally in HIV-1 seropositive individuals (25–26) and the HIV-1 Tg rat (26). Indeed, older adult HIV-1 Tg rodents, independent of sex, exhibit increased Aβ protein levels relative to control animals (Main Effect: Genotype, [*F*(1,44)=7.9, *p*≤0.001]; Figure 4A-C).

**Figure 4.**
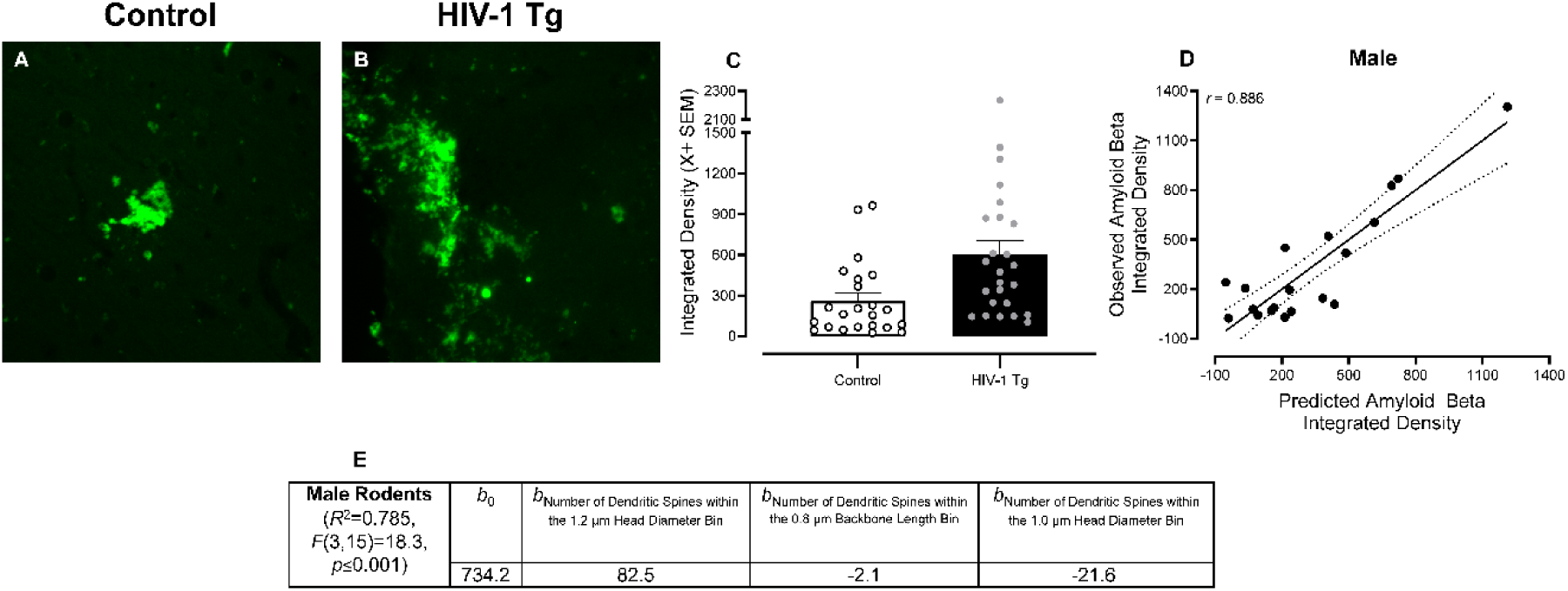
Amyloid Beta (Aβ) Abnormally Accumulates in Rats with Constitutive Expression of HIV-1 Viral Proteins. **(A-B)** Fluorescent immunohistochemical approaches were implemented to evaluate Aβ accumulation in the medial prefrontal cortex of HIV-1 Tg and control animals. **(C)** Quantification of fluorescent Aβ signals revealed increased Aβ protein levels in older adult HIV-1 Tg rodents relative to controls. **(D-E)** A multiple linear regression approach was implemented to evaluate if and/or how Aβ protein levels are associated with dendritic spine morphology. In male HIV-1 Tg rats 78.6% of the variance in the integrated density of Aβ was accounted for by three variables. As such, multiple linear regression analyses support Aβ as a key neural mechanism underlying synaptodendritic dysmorphology in male HIV-1 Tg rats.

Fundamentally, Aβ protein levels are associated with dendritic spine dysmorphology in male (Figure 4D-E), but not female (Data Not Shown), rodents. Specifically, in male animals, 78.6% of the variance in the integrated density of Aβ was accounted for by three variables, including the number of dendritic spines within the 1.0 μm and 1.2 μm head diameter bin and the number of dendritic spines within the 0.8 μm backbone length bin ([*F*(3,15)=18.3, *p*≤0.001]).

### Exploratory multiple regression analyses support neuronal dysfunction as a putative neural mechanism underlying neurocognitive impairments in older adult rodents

An exploratory discriminant function analysis was utilized to determine which neurocognitive assessments were best able to identify genotype. A stepwise discriminant function analysis selective three variables (Visual PPI Area Under the Curve, Number of Days Meeting Criteria in the Visual Distractor Task, and Number of Hits During Block 2 of the Visual Distractor Task) that maximally separated the HIV-1 Tg and control animals (Figure 5A; Canonical Correlation of 0.661). Rodents were classified (jackknifed) with 84.1% accuracy (*F* approximation of Wilks’ λ of 0.563, [*F*(3,42)=10.4, *p*≤0.001]).

**Figure 5.**
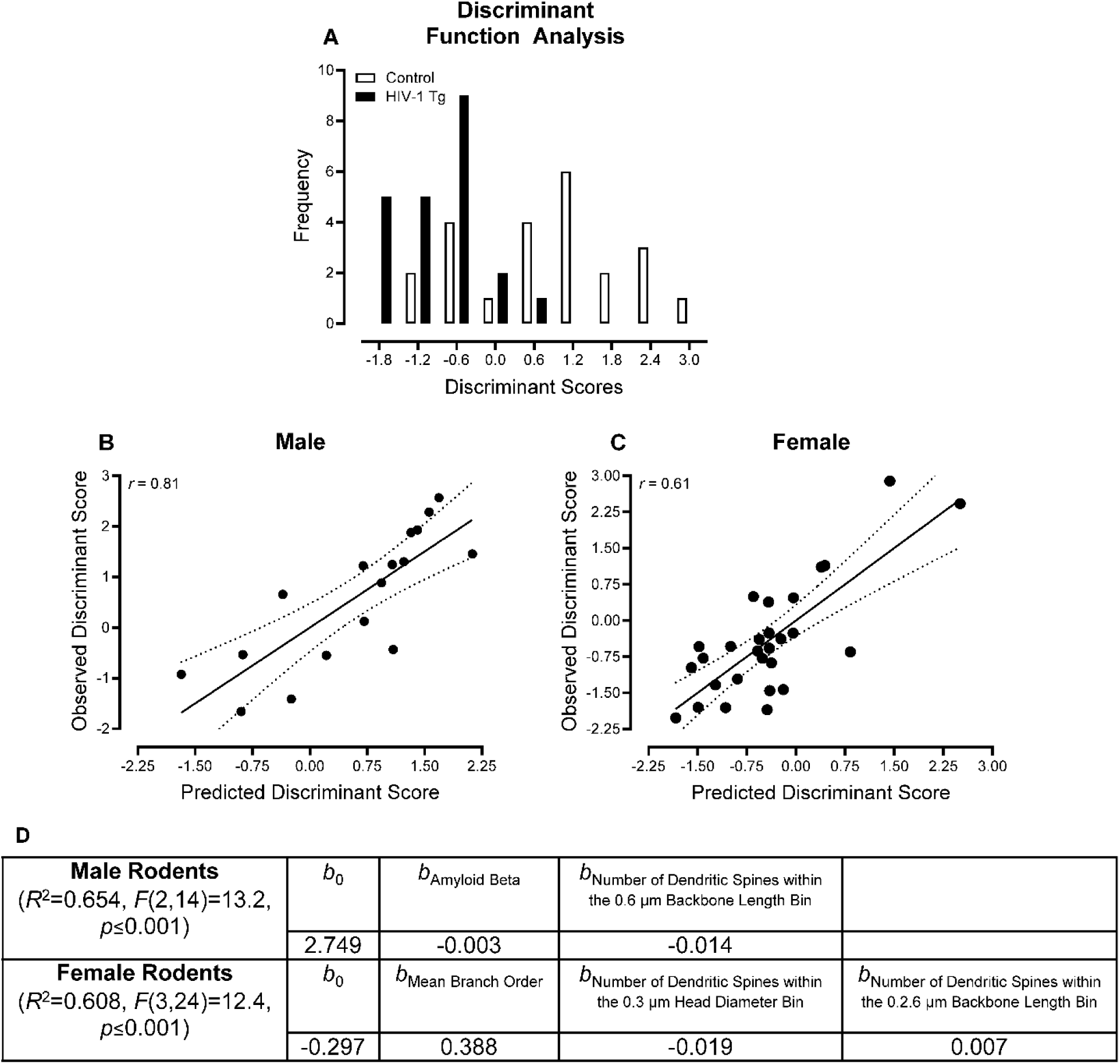
Neuronal Dysfunction Mechanistically Underlies, At Least in Part, Neurocognitive Impairments in Older Adult HIV-1 Transgenic (Tg) Rats. **(A)** An exploratory discriminant function analysis revealed three variables, including Visual PPI Area Under the Curve, Number of Days Meeting Criteria in the Visual Distractor Task, and Number of Hits During Block 2 of the Visual Distractor Task) that maximally separated HIV-1 Tg and control animals. Rodents were classified (jackknifed) with 84.1% accuracy. **(B-D)** A multiple linear regression approach was implemented to evaluate the relationship between neurocognitive function and neuronal dysfunction (e.g., Aβ Integrated Density, Mean Branch Order, Dendritic Spine Backbone Length and Head Diameter). Neurocognitive function was indexed using a discriminant score, which was calculated for each rodent based on the discriminant function analysis. In male HIV-1 Tg rats, 65.4% of the variance in neurocognitive function was accounted for by two variables. In female HIV-1 Tg rodents, 60.8% of the variance in neurocognitive function was accounted for by three variables.

For each rodent, a discriminant score was calculated by the statistical software and utilized for subsequent multiple regression analyses. Indeed, the multiple regression analysis was conducted to evaluate the relationship between neurocognitive function, indexed by the discriminant score, and neuronal dysfunction (e.g., Aβ Integrated Density, Mean Branch Order, Dendritic Spine Backbone Length and Head Diameter). In male rodents, 65.4% of the variance in neurocognitive function was accounted for by two variables, including the Aβ integrated density and the number of dendritic spines within the 0.6 μm backbone length bin (Figure 5B,D; [*F*(2,14)=13.2, *p*≤0.001]). In female animals, 60.8% of the variance in neurocognitive function was accounted for by three variables (i.e., Mean Branch Order, Number of Dendritic Spines within the 0.3 μm Head Diameter Bin and 2.6 μm Backbone Length Bin; Figure 5C,D; [*F*(3,24)=12.4, *p*≤0.001]). Exploratory multiple regression analyses, therefore, support neuronal damage as a key mechanism underlying neurocognitive dysfunction.

## DISCUSSION

The population of older adult HIV-1 Tg rodents sampled exhibited profound neurocognitive impairments at least partly resulting from neuronal dysfunction in the mPFC. Using a longitudinal experimental design without the confound of repeated testing, the true magnitude of neurocognitive and neuroanatomical decline in the HIV-1 Tg rat was evaluated. Consistent with prior reports (16), older adult HIV-1 Tg and control rodents exhibited intact visual and gross-motoric system function; findings which support the utility of the HIV-1 Tg rat for studies investigating age-related neurocognitive impairments. Indeed, the population of older adult HIV-1 Tg rodents sampled displayed profound impairments in select neurocognitive domains, including stimulus-reinforcement learning, sustained attention, and selective attention. Visual discrimination and extradimensional set-shifting, in sharp contrast, remained relatively spared. Prominent neuronal dysfunction was characterized by structural alterations in the morphological parameters of pyramidal neurons, and their associated dendritic spines, and exacerbated Aβ accumulation in the mPFC. Fundamentally, neuronal dysfunction accounted for 60.8% to 65.4% of the variance in neurocognitive impairments in older adult HIV-1 Tg and control female and male animals, respectively. Thus, without the confound of repeated testing, the true magnitude of neurocognitive and neuroanatomical decline in the HIV-1 Tg rat was revealed supporting a biological system to model age-related neurocognitive impairments.

Older adult HIV-1 Tg rodents exhibited a fundamental distortion of temporal processing evidenced by selective neurocognitive impairments in stimulus-reinforcement learning, sustained attention, and selective attention. Temporal processing, which is deeply integrated with higher-order cognitive processes (e.g., attention, memory), describes the brain’s ability to process and interpret successive stimuli across time. Indeed, the systematic manipulation of stimulus duration (i.e., in the signal detection operant task or visual discrimination task) placed significant demands on temporal processing in the HIV-1 Tg rat resulting in deficits in sustained and selective attention. Temporal processing at longer intervals (i.e., days) also appears dysregulated, as stimulus-reinforcement learning was impaired in older adult HIV-1 Tg rodents, evidenced by the increased number of days required to meet criteria. Distorted timing appears to be a critical component of HAND, as temporal processing deficits have been observed during multiple developmental periods (e.g., Preweanling/Childhood: (27); Adulthood: (28); Older Adults: Present Study) and progress across time in the HIV-1 Tg rat (16–17); the subjective experience of time’s passage (i.e., time perception), which relies upon intact temporal processing, is also impaired in HIV-1 seropositive individuals (29–30).

The primary function of the prefrontal cortex (PFC), an anatomically complex brain region that receives projections from thalamic nuclei (31), is to manage higher-order cognitive processes (i.e., executive functions), including attention, flexibility, and cognitive control. Structurally, the agranular PFC is organized in a laminar fashion and is comprised of three major subdivisions, including the mPFC, orbitofrontal cortex (OFC), and lateral PFC (lPFC). The mPFC, specifically, is crucial for temporal processing, as evidenced by lesion studies in rodents, primates, and humans (reviewed by (32)). Similarly, individuals with neurological diseases damaging the mPFC, including schizophrenia, HIV-1, and substance use disorders, experience significant impairments in temporal processing (Schizophrenia: (33); HIV-1: (29–30); Substance Use Disorders: (34–35)). Recent technological advances have further afforded scientists an opportunity to identify the neural circuits involved in temporal processing, whereby both the striatum (36) and cerebellum (37) interact with the mPFC to encode temporal predictions. The structural basis of neural circuits is formed by dendritic spines, which are morphologically heterogeneous protrusions emanating from the dendritic shaft of a neuron. The highly dynamic nature of dendritic spine morphology is of fundamental importance to neural circuit plasticity, as the morphological parameters of dendritic spines shift in response to experience (e.g., Learning and Memory Formation; (38)) and pathological conditions (e.g., Schizophrenia: (39); HIV-1: (40–42)). Classically, dendritic spines are categorized into one of four distinct groups (i.e., filopodia, thin, mushroom, stubby) based on their morphological parameters, whereby long, thin filopodia and strong, stable mushroom spines represent the most immature and mature dendritic spines, respectively. Nevertheless, utilization of a reductionist approach (i.e., classification of dendritic spines) results in the loss of nuanced information and precludes inferences regarding synaptic function (43). Indeed, there is a strong positive association (*r* = 0.88) between the size of the dendritic spine head and area of the postsynaptic density (PSD; (44–45)); dendritic spine backbone length is inversely correlated (*r* = - 0.72) with excitatory postsynaptic potentials (46). Herein, investigation of dendritic spine morphology along a continuum revealed a prominent sex- and genotype-dependent population shift in dendritic spine backbone length and head diameter supporting HIV-1-induced alterations in synaptic function resulting in the dysregulation of neural circuits.

Accumulation of Aβ mechanistically underlies, at least in part, the profound dendritic spine dysmorphology in male, but not female, HIV-1 Tg rodents. Aβ peptides are formed following the proteolysis of the amyloid precursor protein along the amyloidogenic, rather than the nonamyloidogenic, pathway. Classically, insoluble extracellular Aβ plaques formed by the aggregation of Aβ peptides are considered a hallmark late-stage neuropathological feature of Alzheimer’s disease (47–48); intraneuronal accumulation of Aβ is associated with early Alzheimer’s disease damage (49–51). In sharp contrast, Aβ peptide accumulation in HIV-1 seropositive individuals (25) and the HIV-1 Tg rat (26) occurs predominantly within neurons. Mechanistically, the HIV-1-induced abnormal intraneuronal accumulation of Aβ likely results from alterations in Aβ synthesis and/or Aβ degradation (25, 52–54). Fundamentally, Aβ impairs synaptic plasticity, evidenced by structural alterations in the morphological parameters of dendritic spines (55), disruption of the PSD (for review, 56), impairment of long-term potentiation (57–58), and enhancement of long-term depression (59). Indeed, the implementation of multiple linear regression techniques confirmed and expanded upon prior literature, whereby Aβ integrated density accounted for 78.5% of the variance in dendritic spine morphology in male HIV-1 Tg rodents.

Advanced statistical approaches (i.e., discriminant function analyses, multiple linear regression techniques) also afforded a critical opportunity to establish neuronal dysfunction as a key mechanism underlying neurocognitive impairments in older adult HIV-1 Tg rats. At the structural level, synaptodendritic dysfunction in HIV-1 seropositive individuals during the post-cART era is characterized by neuronal injury (60) and significant reductions in synaptic markers/density (61–63); structural alterations that have been recapitulated and extended in multiple biological systems utilized to model key aspects of HIV-1 (e.g., Simian Immunodeficiency Virus: (62); Transactivator of Transcription (Tat) Tg mice: (40); gp120 Tg mice: (64); Chimeric HIV Rat: (65)). In the HIV-1 Tg rat, specifically, the profound structural alterations in neurons and their associated dendritic spines generalize across multiple brain region (e.g., PFC: (17, 42); Nucleus Accumbens: (41)) and progress across time (66). With regards to neuronal function, the HIV-1 viral protein Tat induces region-specific effects on the electrophysiological properties of neurons characterized, in part, by an increase in excitability in the mPFC (67). Fundamentally, in the post-cART era, structural and functional alterations have been strongly associated with neurocognitive impairments in HIV-1 seropositive individuals (*r* = -0.575 - -0.734 in Frontal Cortex; (68)) and the HIV-1 Tg rat (*r* = 0.78 – 0.81; Present Study).

Although there are several biological systems available to model key aspects of HIV-1, to date, there is no suitable extant *in vivo* biological system to address scientific questions at the intersection of HIV-1 and aging (12). In a longitudinal study, the utility of the HIV-1 Tg rat for assessing the progression of neurocognitive impairment was established (16–17); confounds created by repeated testing, however, may have resulted in an underestimation of the rate of neurocognitive decline induced by constitutive expression of HIV-1 viral proteins. Utilization of a longitudinal experimental design without repeated testing, as in the present study, revealed the true magnitude of neurocognitive and neuroanatomical decline in older adult HIV-1 Tg rodents relative to their control counterparts. Nevertheless, the constitutive expression of HIV-1 viral proteins throughout development in the HIV-1 Tg rat (14–15) precludes differentiation of aging and duration of infection. As such, there remains a fundamental need to critically evaluate the utility of other biological systems (e.g., Simian Immunodeficiency Virus, Chimeric HIV-1, Humanized Mice) for establishing if and/or how duration of infection influences neurocognitive decline during the aging process.

Collectively, a longitudinal experimental design without the confound of repeated testing revealed the true magnitude of neurocognitive and neuroanatomical decline in the HIV-1 Tg rat. Indeed, the population of older adult HIV-1 Tg rodents sampled exhibited profound, albeit selective, neurocognitive impairments characterized by a fundamental distortion of temporal processing. From a mechanistic perspective, neurocognitive impairments resulted, at least partly, from an abnormal intraneuronal Aβ accumulation and dendritic spine dysmorphology. Establishing the utility of the HIV-1 Tg rat for age-related neurocognitive impairments is fundamental to disentangling the role of HIV-1 viral proteins and comorbidities in neurocognitive function.

## ACKNOWLEDGEMENTS/AUTHOR CONTRIBUTIONS

All authors have accepted responsibility for the entire content of this manuscript and approved its submission. Conceptualization, C.F.M. and R.M.B.; Data Collection, K.A.M., A.R., H.L.; Data Analysis, K.A.M., A.R., and C.F.M.; Writing—Original Draft Preparation, K.A.M.; Review and Editing, K.A.M., R.M.B., and C.F.M.; Funding Acquisition, C.F.M., R.M.B., and K.A.M. All authors have read and agreed to the published version of the manuscript.

## DATA AVAILABILITY STATEMENT

The authors declare that all the data supporting the findings of this study are available within the paper and its Supplemental Data.

