## Supplementary file for "IMPAIRED BRIDGING OF TEMPORAL DISCONTINUITIES IN OLDER ADULT HIV-1 TG RATS"

### IMPAIRED BRIDGING OF TEMPORAL DISCONTINUITIES IN THE AGING HIV-1 TG RAT

#### MATERIALS AND METHODS

##### Integrity of Visual System Function: Visual Prepulse Inhibition.

***Procedure***. Prior to the cross-modal PPI assessment, rats (Control Male: *n*=12; Control Female: *n*=16; HIV-1 Tg Male: *n*=10; HIV-1 Tg Female: *n*=17) completed a habituation session using the methodology outlined by McLaurin et al. (1). The PPI test session began with a five-minute adjustment period followed by six pulse-only auditory startle response (ASR) trials (Intertrial Interval: 10 sec). Subsequently, a counterbalanced experimental design (i.e., ABBA) was utilized for the presentation of 72 testing trials (Intertrial Interval: 15-25 sec), which included an equal number of auditory and visual prepulse trials. During the testing trials, the time between the prepulse and startle stimulus (i.e., Interstimulus Interval; 0, 30, 50, 100, 200 or 4000 msec) was systematically manipulated, whereby ISIs were presented in 6-trial blocks according to a Latin Square experimental design. Peak ASR amplitude values were collected for analysis.

##### Components of Executive Function

***Sustained Attention: Signal Detection Operant Task.*** A response on the correct lever during a signal trial (i.e., central panel light illumination) defined as a hit, whereas a correct response during a non-signal trial (i.e., no illumination) is categorized as a correct rejection; both response types are indicative of attention to the stimulus. Responding on the incorrect lever during a signal or non-signal trial is defined as a miss or false alarm, respectively, whereby a miss represents a lapse of attention, and a false alarm supports a failure of response inhibition. Rodents were required to respond at least five times and achieve 65% accuracy, calculated as (((Number of Hits and Correct Rejections at 1000 msec)/(Total Number of Responses at 1000 msec)) x 100) for three consecutive or five non-consecutive days to be promoted to the next vigilance program. The methods utilized to train animals in the signal detection operant task are described in detail by McLaurin et al. (2). Statistical analyses and figures represent the days meeting criteria during the third vigilance program, yielding samples sizes of: (Control Male: *n*=12; Control Female: *n*=14; HIV-1 Tg Male: *n*=7; HIV-1 Tg Female: *n*=17).

**Post-mortem Analysis**

***Synaptodendritic Dysfunction: Ballistic Labeling.*** Ballistic cartridges were created by drawing a tungsten bead/DiIC19(3) dye mixture into polyvinylpyrrolidone-coated Tefzel tubing, drying the mixture with nitrogen gas (0.5 Liters per Minute) for 30 minutes, and cutting the tubing into 13 mm lengths. The prepared ballistic cartridges were loaded into the Helios gene gun (Bio-Rad, Hercules, CA, USA), which was connected to a helium hose (Output Pressure: 90 pounds per square inch), and the applicator was used to fire the tungsten bead/DiIC19(3) bead tubing onto the brain slices. Tissue slices were washed three times with 100 mM PBS, stored in the dark at 4ºC for three hours, and mounted onto a glass slide using Pro-Long Gold Antifade (Invitrogen, Carlsbad, CA, USA).

Z-stack images (60x Oil Objective, *n.a.*=1.4, Z-Plane: 0.15 µm) of pyramidal neurons in the mPFC were acquired using a Nikon TE-2000E confocal microscopy system in combination with Nikon's EZ-C1 software (Version 3.81b).

Pyramidal neurons, and associated dendritic spines, from the mPFC were analyzed using Neurolucida 360 (MicroBrightfield, Williston, VT, USA). After blinding, stringent selection criteria (e.g., low background/dye clusters, continuous dendritic staining; (3)) were implemented to select one neuron from each animal, yielding the following sample sizes: (Control Male: *n*=12; Control Female: *n*=11; HIV-1 Tg Male: *n*=7; HIV-1 Tg Female: *n*=18).

The classic Sholl analysis and a centrifugal branch ordering scheme were utilized to evaluate key aspects of neuronal morphology, including dendritic arbor complexity and dendritic branching complexity, respectively. Boundary conditions for dendritic spines were established (Volume: 0.05 to 0.85 µm; Backbone Length: 0.4 to 4.0 µm; Head Diameter: 0.001 to 1.2 µm), whereby only dendritic spines meeting all of the boundary conditions were included in the analysis. Two parameters, including dendritic spine backbone length and head diameter, were analyzed to evaluate dendritic spine morphology.

**Statistical Analysis**

The integrity of visual (Visual PPI) and gross motoric (Locomotor Activity) system function, as well as amyloid beta fluorescent signals, were statistically evaluated using a repeated-measures ANOVA (SPSS Statistics 31; Visual PPI) or univariate ANOVA (SPSS Statistics 31; Locomotor Activity, Amyloid Beta). Specifically, genotype (HIV-1 Tg vs. Control) and biological sex (Male vs. Female) served as between-subjects factors, whereas ISI served as a within-subjects factor, as appropriate. For gross motoric system function, complementary one sample *t*-tests were conducted independently for each genotype to statistically test whether the average number of photocell interruptions was significantly greater than 0.

Generalized linear mixed models (PROC GLIMMIX, SAS/STAT Software 9.4) were utilized to statistically evaluate the animals' response profile during neurocognitive assessments (i.e., Sustained Attention, Selective Attention, Discrimination, Extradimensional Set-Shifting), the number of hits and misses across signal durations during neurocognitive tasks tapping sustained or selective attention, and neuronal (i.e., Branch Order) and dendritic spine dysmorphology. The model was calculated using a Poisson distribution with a log link, a random intercept, and either an AR(1) or compound symmetry covariance structure (AR(1): Neurocognitive Assessments; Compound Symmetry: Neuroanatomical Assessments). Analyses for sustained attention, selective attention, and branch order were conducted on data transformed by squaring the original values. For analyses of selective attention, rodents with fewer than an average of two responses during the second trial block (i.e., when the visual distractor was present) were censored. The factor of biological sex was included in analyses of sustained attention, as well as neuronal and dendritic spine dysmorphology, but not in analyses of selective attention, discrimination, or extradimensional set-shifting.

The Sholl intersection profile was analyzed using a mixed-model ANOVA with a variance components covariance structure (PROC MIXED; SAS/STAT Software 9.4; (4)).

SPSS Statistics 31 was utilized to conduct an exploratory discriminant function analysis and multiple linear regression analyses. Four rodents (HIV-1 Tg Male: $n$=3; HIV-1 Tg Female: $n$=1) were not included in the exploratory discriminant function analysis as they were missing at least one discriminating variable.

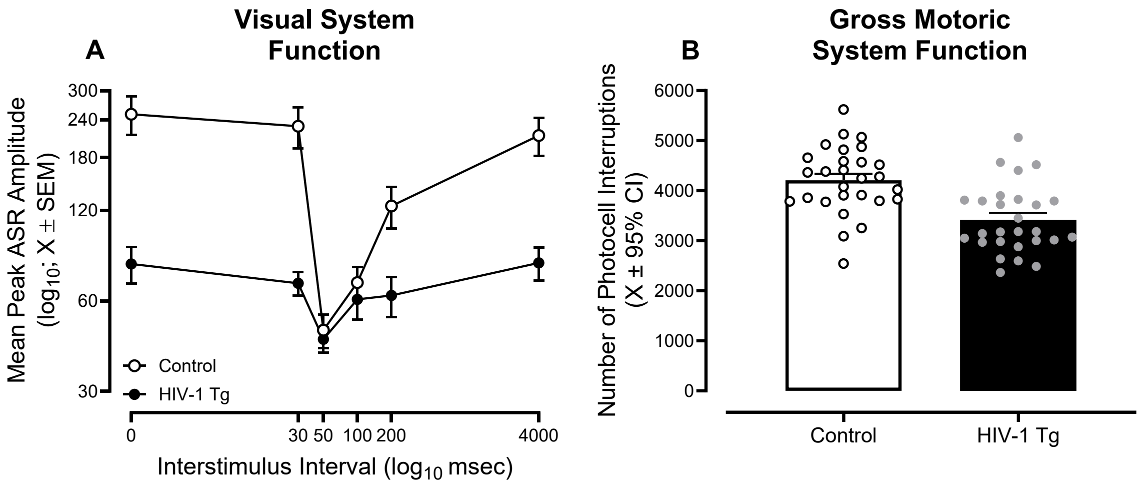


**Supplementary Figure 1. HIV-1 Transgenic (Tg) Rats Exhibit Intact Visual and Gross Motoric System Function. (A)** Visual prepulse inhibition of the auditory startle response was conducted to evaluate the integrity of visual system function in older HIV-1 Tg and control animals. Independent of genotype, robust inhibition to the presence of a visual prepulse at the 50 msec interstimulus interval supports the integrity of visual system function. **(B)** Rodents were evaluated in three 60-min locomotor activity tests sessions, whereby the average number of photocell interruptions across all three test sessions is presented. Although HIV-1 Tg rodents exhibited a significantly fewer mean number of photocell interruptions relative to control animals, the average number of photocell interruptions were significantly greater than 0.
